# Imidazole-imidazole hydrogen bonding in the pH sensing Histidine sidechains of Influenza A M2

**DOI:** 10.1101/802942

**Authors:** Kumar Tekwani Movellan, Melanie Wegstroth, Kerstin Overkamp, Andrei Leonov, Stefan Becker, Loren B. Andreas

**Affiliations:** Department of NMR Based Structural Biology, Max Planck Institute for Biophysical Chemistry, Am Fassberg 11 Göttingen, Germany

## Abstract

The arrangement of histidine sidechains in influenza A M2 tetramer determines their pKa values, which define pH controlled proton conduction critical to the virus lifecycle. Both water associated and hydrogen bonded Imidazole–Imidazolium histidine quaternary structures have been proposed, based on crystal structures, and NMR chemical shifts, respectively. Here we show, using the conduction domain construct of M2 in lipid bilayers, that the imidazole rings are hydrogen bonded even at a pH of 7.8 in the neutral charge state.

An intermolecular 8.9 ± 0.3 Hz ^2h^J_NN_ hydrogen bond is observed between H37 N_ε_ and N_δ_ recorded in a fully protonated sample with 100 kHz magic-angle spinning. This interaction could not be detected in the drug-bound sample.

---

The M2 protein from Influenza A assembles as a tetramer^1-3^ and is tuned to conduct protons across the virus envelope upon external acidification during endocytosis, which leads to membrane fusion and release of RNA into the host organism.^3-4^ This process is controlled by the quaternary structure of the tetramer and the protonation state of the four pH sensitive H37 residues. Reported pKa values for H37 range from 6.3 to 8.2, for the first two protonation events, with a consensus that lies above the value of 6 found for the histidine side-chain in aqueous solution.^5-8^ Correlating these pKa measurements with the endosomal pH resulted in the understanding that the 3rd proton to enter the channel results in conduction.^7^ Yet despite many reports of the M2 structure from oriented sample NMR,^9-11^ solution NMR,^12-14^ magic-angle spinning NMR^15-17^ and crystallography^18-21^ there is still no consensus in the literature regarding the structural configuration at H37 that leads to these shifted pKa values.

Based on ^15^N chemical shifts, Cross and coworkers proposed that the doubly charged tetramer arranges its histidine sidechains to form imidazole-imidazolium dimers, delocalizing the positive charge and explaining the high pKa.^7^ This imidazole arrangement is not seen in crystal structures,^18-21^ either at high or low pH. Instead, changes in pH are associated to opening or closing of the c-terminal side of the tetramer, rather than changes in the geometry of the histidine side chain. In a recent high resolution x-ray free electron laser (XFEL) crystal structure, the histidine is found hydrogen bonded to water with H37 residues separated ∼7 Å apart, too far to form a hydrogen bond.^21^

In further support of the hydrogen bonded dimer arrangement, Cross and coworkers measured proton shifts up to 18.5 ppm for a full-length M2 sample at pH 6.2 where the +2 charge state (half imidazole, half imidazolium) is expected to be the dominant form.^8^ These results support the presence of a strong imidazole-imidazolium hydrogen bond, since strong hydrogen bonds with a low barrier are associated with proton chemical shifts above 16 ppm.^22-24^ On the other hand, Hong and coworkers recorded HETCOR measurements in the transmembrane (TM) construct (residues 22-46) that showed strong contacts to water and proton shifts between 8 and 15 ppm, consistent with normal hydrogen bonds.^25^

While the XFEL and NMR data strongly support that H37 in the TM construct forms hydrogen bonds to water, so far only indirect evidence in the form of chemical shifts exists in support of the dimer configuration. The 18.5 ppm proton shift observed in M2 is also not far from a proton shift of 16.8 ppm that was seen for H_δ1_ in an N-H- -O hydrogen bond found in the free amino acid histidine^26^ and therefore does not unequivocally identify the hydrogen bonding partner. Although indirect, this evidence was recorded for a full length construct in lipid bilayers, which should more closely match physiological conditions. It is clear from the existing data that the channel is hydrated, yet a direct measurement is needed to prove that a hydrogen bonded histidine dimer is also one of the stable arrangements of the tetramer.

Measurement of J-couplings in NMR is an established method to identify hydrogen bonding interactions. For example, detection of amide nitrogen to carbonyl J-coupling (^3h^J_NC_) establishes hydrogen bonding in proteins both in solution^27-28^ and in microcrystals^29^. The base pairing interactions in DNA and RNA result in N-H- -N hy-drogen bonds and 6-7 Hz J-couplings.^30-31^ An unusual example of a histidine–histidine ^2h^J_NN_ coupling of the type proposed in M2 occurs for the protein apomyoglobin.^32^ A coupling strength of 10 Hz was reported and it was noted that the N-N distance of 2.75 Å in the holomyoglobin crystal structure (PDB 1MBD) is shorter than those of the base pairs (∼2.9 Å). Though these differences are within the error of the measurements, they are in line with the understanding that closer N-N distances result in stronger couplings.

In the ‘conductance domain’ (residues 18-60) M2 tetramer, a C_2_ symmetric dimer of dimers arrangement^33^ at a relatively high pH of 8.5 results in two H37 H_ε2_ chemical shifts, at 12.1 and 14.5 ppm, both in the tau tauto-mer^5^. See the Supplemental Information for the ^1^H-^15^N correlation spectrum recorded at pH 7.8. The question remains whether the fourfold symmetry is broken by hydrogen bonded dimers or whether only water is the hydrogen bonding partner. Although the H37 proton chemical shifts are indicative of normal hydrogen bonds, a dimer arrangement in the neutral charge state would be expected to persist in the +2 state, since the dimer further stabilizes a positive charge, according to *ab initio* quantum chemical calculations on imidazole dimers^34^ and in the core of the M2 tetramer.^35^ Such calculations show that a positive charge leads towards a stronger dimer interaction that might lead to a strong or low barrier hydrogen bond.

We measured an imidazole-imidazole ^2h^J_NN_-coupling using a homonuclear INEPT^36^ period on the nitrogen channel, combined with cross polarization (CP) for detection of the attached proton. (Fig 1) If no J coupling is present, only the peak from the CP based NH spectrum remains. If a homonuclear J coupling is present, then an additional peak is observed at the ^15^N frequency of the coupled spin, and with a buildup of intensity following the well-known relation, sin^2^(2πJt)exp(-τ/t_2_). Since the original peak decays with cos^2^(2πJt)exp(-τ/t_2_), normalization by the total signal at each mixing time results in a single parameter fit to the equation sin^2^(2πJt). We measured an N_δ1_-N_ε2_ J-coupling of 8.9 ± 0.3 Hz Hz (Fig 2), and the frequency indicates a J-coupling to the neighboring inequivalent histidine. The histidine ^15^N and ^1^H assignments were reported previously in Colvin et al.^5^ Consistent with the understanding that the ^1^H shift is correlated with the strength of the hydrogen bonding interaction, this intermolecular N-H- -N ^2h^J_NN_ coupling occurs for the most strongly downfield shifted proton at 14.5 ppm. No homonuclear J-coupling could be detected for the other N_ε2_, indicating that its attached H_ε2_ at 12.1 ppm is likely hydrogen bonded to oxygen.

**Figure 1.**
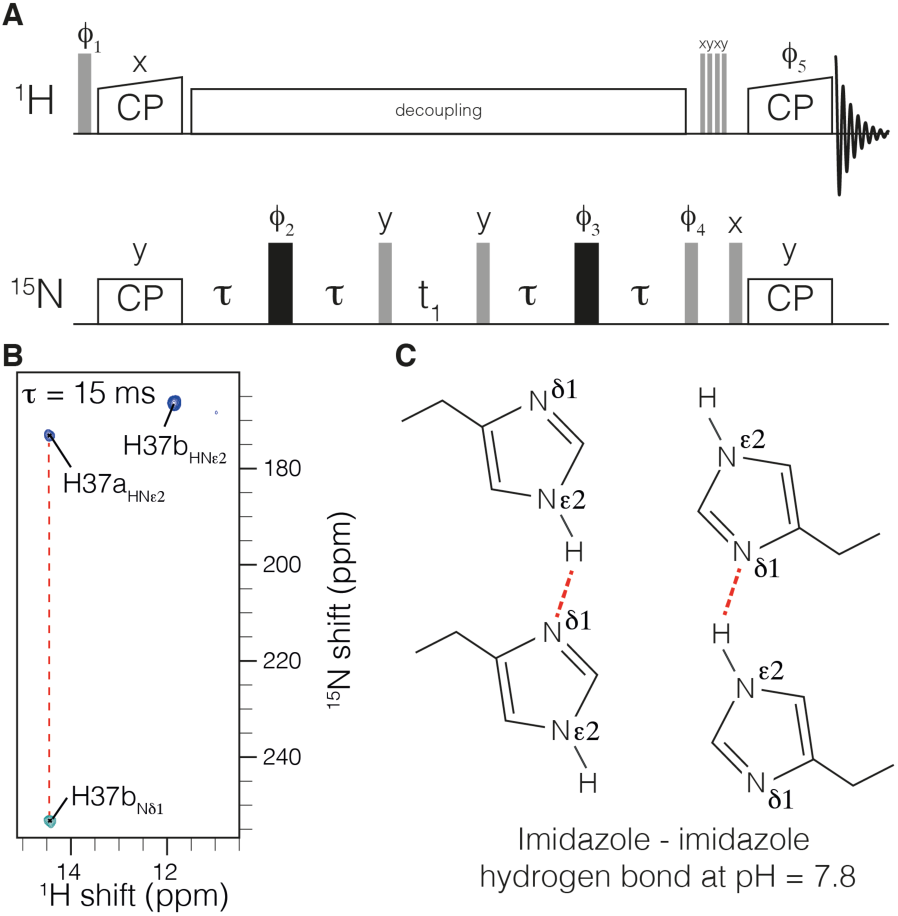
Measurement of ^2h^J_NHN_ hydrogen bonding in H37 imidazole dimers within influenza M2. The pulse sequence is shown in a). Cross-polarization (CP) is used to establish ^15^N polarization. A homonuclear out-and-back INEPT period is then used to record the chemical shift of the J-coupled nitrogen. Following water suppression, CP is used to transfer back to protons for detection. The spectrum in b) was recorded with τ of 15 ms, and clearly shows a negative peak indicative of an intermolecular J-coupling, and a C_2_ symmetric tetramer at H37, as shown schematically in c). The phase cycle was ϕ_1_=y,y,-y,-y ϕ_2_=8·{x},8·{y} ϕ_3_=16·{x},16·{y} ϕ_4_=x,-x ϕ_5_=4·{y},4·{-y} ϕ_rec_={y,-y,-y,y} 2·{-y,y,y,-y} {y,-y,-y,y} {-y,y,y,-y} 2·{y,-y,-y,y} {-y,y,y,-y}.

**Figure 2.**
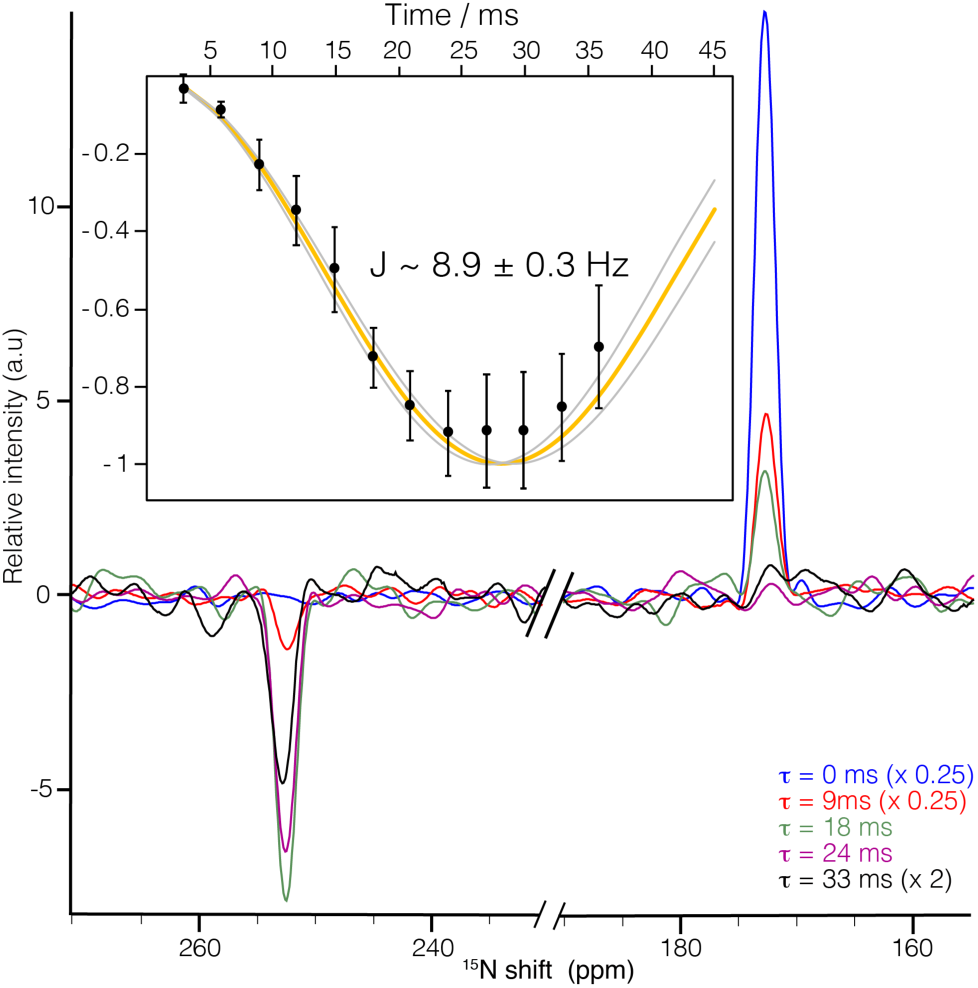
Quantification of the intermolecular ^2h^J_NHN_ J-coupling. Slices of the 2D spectrum at the proton frequency of 14.5 are shown for the indicated mixing times. In the inset, the experimental data (points) were fit as a function of the time τ. Relaxation was accounted for by dividing each intensity at 254 ppm by the total signal magnitude of the slice. The best fit (orange) resulted in a coupling strength of 8.9 ± 0.3 Hz. The curves in grey indicate the error at twice the standard deviation, σ, as estimated with a Monte Carlo approach taking into consideration the random spectral noise. The spectral noise level in the experimental data is displayed at 2σ. The first point was acquired with 8 scans (1.5 hours) while the last point required 128 scans (26 hours) due to transverse relaxation (See Fig. S2).

That both water-associated and hydrogen bonded dimer conformations of H37 exist under different conditions suggests that these conformations are both relatively stable. The main differences in sample preparation that result in these different structures are the nature of the membrane mimetic, and the length of the construct. The fact that the N—H- -N J coupling is observed in the longer ‘conductance domain’ construct embedded in lipid bilayers suggests that it is a functionally relevant state, and that the proton affinities of this state control the interconversion to a conducting channel. (Recall that the third proton is thought to result in conductance, and this third protonation event necessarily breaks the ^2h^J_NN_ hydrogen bond). This does not rule out the possibility that other quaternary structures may also lead to a proton current in virus particles, although a different pKa would be expected at H37. This is in line with the large range of pKa values reported for M2. ^5-8^

Interestingly, in a sample with the inhibitor rimantadine bound to the pore, the histidine H_ε2_ peaks shift upfield by about 3 ppm, and no hydrogen bond could be detected (Fig. 3). This disruption of the hydrogen bonding interaction explains the difference in H37 pKa in the drug bound state,^37^ and lends support to the hypothesis that imidazole dimers are functionally important. Both rimantadine and amantadine bind to the pore of M2, with the simplest explanation for their efficacy being that they plug the pore, preventing passage of hydronium ion. Yet significant chemical shift perturbations^33, 38^ were detected widely over the transmembrane residues. This suggested that the inhibitors have far reaching effects, modifying the conformational distribution of the protein. However, it has been difficult to connect a specific structural change to the chemical shift perturbations. It is now clear that in the conductance domain construct, rimantadine affects the structure of the channel by impacting the hydrogen bonding of the important functional residue H37.

**Figure 3.**
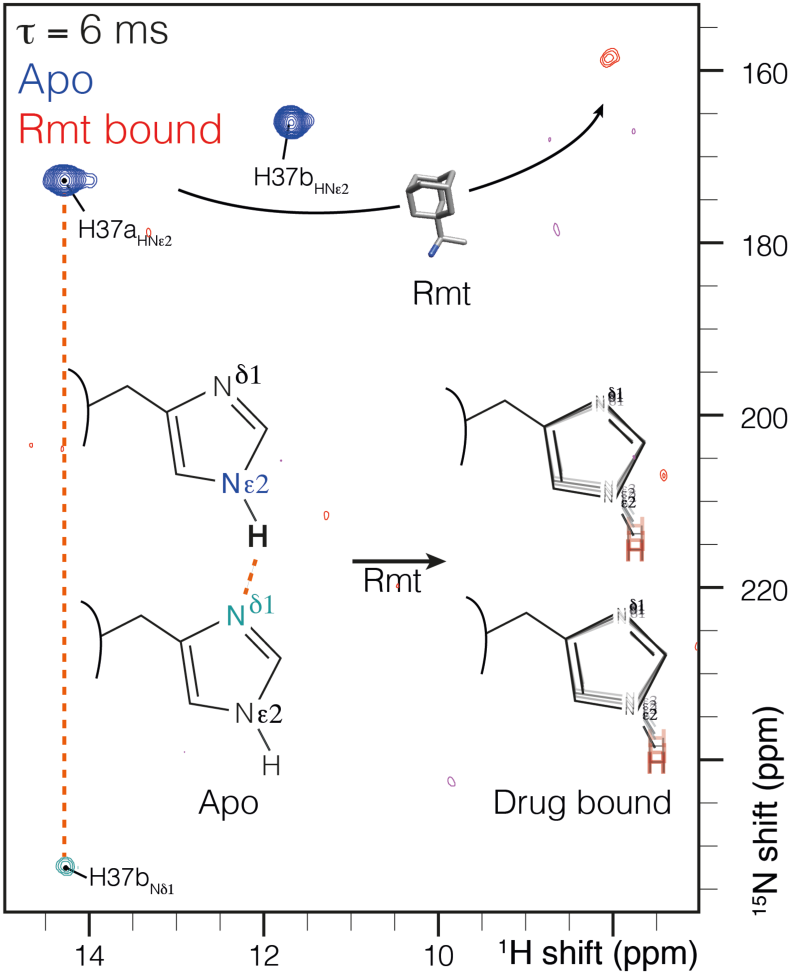
Chemical shift changes in the histidine side-chain upon addition of the drug rimantadine (Rmt) using the pulse sequence of Fig. 1. A 3-5 ppm change is observed in the drug bound spectrum (red). No ^2h^J_NHN_ J-coupling was observed in the drug bound sample. Instead, the imidazole NH peaks are broadened, indicating motion or chemical exchange.

We used the M2_18-60_ construct of M2 in 1,2-diphytanoylsn-*glycero*-3-hosphatidylcholine (DPhPC) bilayers, as in previous work.^33^ This construct has been dubbed the ‘conductance domain’ since it has been shown to recapitulate the proton conduction rates and drug sensitivity properties of the full length protein in liposome flux assays.^12^ The protein was expressed as a fusion to TrpLE, cleaved with cyanogen bromide, and purified by HPLC, as described previously.^13^ The serine at position 50 was changed from the native cysteine. The sequence after cleaving is RSN^20^ DSSDPLVVAA^30^ SIIGILHLIL^40^ WILDRLFFKS^50^ IYRFFEHGLK^60^. Lyophilized protein was resuspended in octyl glucoside detergent and reconstituted in perdeuterated (d78-phytanoyl, d9-choline) DPhPC lipids (FBReagents) at a lipid to protein ratio of 1 by mass. The drug-containing sample was equilibrated with a 40 mM solution of rimantadine after reconstitution. All spectra were acquired on a 950 MHz Bruker Avance III spectrometer using a Bruker 0.7 mm HCDN probe. The spinning frequency and gas flow was set to maintain a sample temperature of about 20 °C, (260 K thermocouple temperature, 500 liters per hour) as determined using the chemical shift of potassium bromide. The proton spectrum was referenced by setting the chemical shift of water to 4.75 ppm. The nitrogen spectrum is reported on the liquid ammonia scale, using the IUPAC relative frequency ratios.

In conclusion, through the measurement of a ^2h^J_NN_ coupling, we confirmed the existence of imidazole– imidazole dimers in the M2 protein from influenza. Such a configuration was proposed to stabilize positive charge in the tetrameric channel. However, direct evidence of the interaction was not previously reported, crystal structures of M2 showed an alternate structure, leading to controversy over whether the dimeric histidine arrangement exists at all in M2. We have solved this controversy though NMR measurements of M2 in lipid bilayers at a relatively high pH where we observe a neutral charge state at the functional H37 residue. The coupling strength is consistent with a normal hydrogen bonding interaction. Binding of the drug rimantadine to the pore of M2 resulted in breaking of this hydrogen bond. It remains to be seen whether evidence can be found that this geometry persists in the important +2 charge state, where imidazole–imidazolium dimers have been proposed, and whether such a state results in a normal hydrogen bond or a low barrier hydrogen bond.

## AUTHOR INFORMATION

### Notes

The authors declare no competing financial interests.

## ACKNOWLEDGMENT

We thank James Chou for providing the expression plasmid. We acknowledge financial support from the Max Planck Society and the DFG Emmy Noether program (grant AN 1316/1-1)

## REFERENCES

1. Kochendoerfer, G. G.; Salom, D.; Lear, J. D.; Wilk-Orescan, R.; Kent, S. B.; DeGrado, W. F., Total chemical synthesis of the integral membrane protein influenza A virus M2: role of its C-terminal domain in tetramer assembly. Biochemistry 1999, 38 (37), 11905–13.

2. Sakaguchi, T.; Tu, Q.; Pinto, L. H.; Lamb, R. A., The active oligomeric state of the minimalistic influenza virus M2 ion channel is a tetramer. Proc. Natl. Acad. Sci. U S A 1997, 94 (10), 5000–5.

3. Sugrue, R. J.; Hay, A. J., Structural characteristics of the M2 protein of influenza A viruses: evidence that it forms a tetrameric channel. Virology 1991, 180 (2), 617–24.

4. Sugrue, R. J.; Bahadur, G.; Zambon, M. C.; Hall-Smith, M.; Douglas, A. R.; Hay, A. J., Specific structural alteration of the influenza haemagglutinin by amantadine. EMBO J. 1990, 9 (11), 3469–76.

5. Colvin, M. T.; Andreas, L. B.; Chou, J. J.; Griffin, R. G., Proton association constants of His 37 in the Influenza-A M218-60 dimer-of-dimers. Biochemistry 2014, 53 (38), 5987–94.

6. Hu, F.; Schmidt-Rohr, K.; Hong, M., NMR detection of pH-dependent histidine-water proton exchange reveals the conduction mechanism of a transmembrane proton channel. J. Am. Chem. Soc. 2012, 134 (8), 3703–13.

7. Hu, J.; Fu, R.; Nishimura, K.; Zhang, L.; Zhou, H. X.; Busath, D. D.; Vijayvergiya, V.; Cross, T. A., Histidines, heart of the hydrogen ion channel from influenza A virus: toward an understanding of conductance and proton selectivity. Proc. Natl. Acad. Sci. U S A 2006, 103 (18), 6865–70.

8. Miao, Y.; Fu, R.; Zhou, H. X.; Cross, T. A., Dynamic Short Hydrogen Bonds in Histidine Tetrad of Full-Length M2 Proton Channel Reveal Tetrameric Structural Heterogeneity and Functional Mechanism. Structure 2015, 23 (12), 2300–2308.

9. Hu, J.; Asbury, T.; Achuthan, S.; Li, C.; Bertram, R.; Quine, J. R.; Fu, R.; Cross, T. A., Backbone structure of the amantadine-blocked trans-membrane domain M2 proton channel from Influenza A virus. Biophys. J. 2007, 92 (12), 4335–43.

10. Sharma, M.; Yi, M.; Dong, H.; Qin, H.; Peterson, E.; Busath, D. D.; Zhou, H. X.; Cross, T. A., Insight into the mechanism of the influenza A proton channel from a structure in a lipid bilayer. Science 2010, 330 (6003), 509–12.

11. Wang, J.; Kim, S.; Kovacs, F.; Cross, T. A., Structure of the transmembrane region of the M2 protein H(+) channel. Protein Sci. 2001, 10 (11), 2241–50.

12. Pielak, R. M.; Schnell, J. R.; Chou, J. J., Mechanism of drug inhibition and drug resistance of influenza A M2 channel. Proc. Natl. Acad. Sci. U S A 2009, 106 (18), 7379–84.

13. Schnell, J. R.; Chou, J. J., Structure and mechanism of the M2 proton channel of influenza A virus. Nature 2008, 451 (7178), 591–5.

14. Pielak, R. M.; Chou, J. J., Solution NMR structure of the V27A drug resistant mutant of influenza A M2 channel. Biochem. Biophys. Res. Commun. 2010, 401 (1), 58–63.

15. Andreas, L. B.; Reese, M.; Eddy, M. T.; Gelev, V.; Ni, Q. Z.; Miller, E. A.; Emsley, L.; Pintacuda, G.; Chou, J. J.; Griffin, R. G., Structure and Mechanism of the Influenza A M218-60 Dimer of Dimers. J. Am. Chem. Soc. 2015, 137 (47), 14877–86.

16. Cady, S. D.; Mishanina, T. V.; Hong, M., Structure of amantadine-bound M2 transmembrane peptide of influenza A in lipid bilayers from magic-angle-spinning solid-state NMR: the role of Ser31 in amantadine binding. J. Mol. Biol. 2009, 385 (4), 1127–41.

17. Cady, S. D.; Schmidt-Rohr, K.; Wang, J.; Soto, C. S.; Degrado, W. F.; Hong, M., Structure of the amantadine binding site of influenza M2 proton channels in lipid bilayers. Nature 2010, 463 (7281), 689–92.

18. Stouffer, A. L.; Acharya, R.; Salom, D.; Levine, A. S.; Di Costanzo, L.; Soto, C. S.; Tereshko, V.; Nanda, V.; Stayrook, S.; DeGrado, W. F., Structural basis for the function and inhibition of an influenza virus proton channel. Nature 2008, 451 (7178), 596–9.

19. Acharya, R.; Carnevale, V.; Fiorin, G.; Levine, B. G.; Polishchuk, A. L.; Balannik, V.; Samish, I.; Lamb, R. A.; Pinto, L. H.; DeGrado, W. F.; Klein, M. L., Structure and mechanism of proton transport through the transmembrane tetrameric M2 protein bundle of the influenza A virus. Proc. Natl. Acad. Sci. U S A 2010, 107 (34), 15075–80.

20. Thomaston, J. L.; Wu, Y.; Polizzi, N.; Liu, L.; Wang, J.; DeGrado, W. F., X-ray Crystal Structure of the Influenza A M2 Proton Channel S31N Mutant in Two Conformational States: An Open and Shut Case. J. Am. Chem. Soc. 2019, 141 (29), 11481–11488.

21. Thomaston, J. L.; Woldeyes, R. A.; Nakane, T.; Yamashita, A.; Tanaka, T.; Koiwai, K.; Brewster, A. S.; Barad, B. A.; Chen, Y.; Lemmin, T.; Uervirojnangkoorn, M.; Arima, T.; Kobayashi, J.; Masuda, T.; Suzuki, M.; Sugahara, M.; Sauter, N. K.; Tanaka, R.; Nureki, O.; Tono, K.; Joti, Y.; Nango, E.; Iwata, S.; Yumoto, F.; Fraser, J. S.; DeGrado, W. F., XFEL structures of the influenza M2 proton channel: Room temperature water networks and insights into proton conduction. Proc. Natl. Acad. Sci. U S A 2017, 114 (51), 13357–13362.

22. Brown, S. P., Applications of high-resolution H-1 solid-state NMR. Solid State Nucl. Mag. 2012, 41, 1–27.

23. Lorente, P.; Shenderovich, I. G.; Golubev, N. S.; Denisov, G. S.; Buntkowsky, G.; Limbach, H. H., H-1/N-15 NMR chemical shielding, dipolar N-15, H-2 coupling and hydrogen bond geometry correlations in a novel series of hydrogen-bonded acid-base complexes of collidine with carboxylic acids. Magn. Reson. Chem. 2001, 39, S18–S29.

24. Sharif, S.; Fogle, E.; Toney, M. D.; Denisov, G. S.; Shenderovich, I. G.; Buntkowsky, G.; Tolstoy, P. M.; Huot, M. C.; Limbach, H. H., NMR localization of protons in critical enzyme hydrogen bonds. J. Am. Chem. Soc. 2007, 129 (31), 9558-+.

25. Hong, M.; Fritzsching, K. J.; Williams, J. K., Hydrogen-bonding partner of the proton-conducting histidine in the influenza M2 proton channel revealed from 1H chemical shifts. J. Am. Chem. Soc. 2012, 134 (36), 14753–5.

26. Li, S.; Hong, M., Protonation, tautomerization, and rotameric structure of histidine: a comprehensive study by magic-angle-spinning solid-state NMR. J. Am. Chem. Soc. 2011, 133 (5), 1534–44.

27. Cornilescu, G.; Hu, J. S.; Bax, A., Identification of the hydrogen bonding network in a protein by scalar couplings. J. Am. Chem. Soc. 1999, 121 (12), 2949–2950.

28. Cordier, F.; Grzesiek, S., Direct observation of hydrogen bonds in proteins by interresidue (3h)J(NC ‘) scalar couplings. J. Am. Chem. Soc. 1999, 121 (7), 1601–1602.

29. Schanda, P.; Huber, M.; Verel, R.; Ernst, M.; Meier, B. H., Direct detection of (3h)J(NC’) hydrogen-bond scalar couplings in proteins by solid-state NMR spectroscopy. Angew. Chem. Int. Ed. Engl. 2009, 48 (49), 9322–5.

30. Dingley, A. J.; Grzesiek, S., Direct observation of hydrogen bonds in nucleic acid base pairs by internucleotide (2)J(NN) couplings. J. Am. Chem. Soc. 1998, 120 (33), 8293–8297.

31. Pervushin, K.; Ono, A.; Fernandez, C.; Szyperski, T.; Kainosho, M.; Wuthrich, K., NMR scalar couplings across Watson-Crick base pair hydrogen bonds in DNA observed by transverse relaxation-optimized spectroscopy. Proc. Natl. Acad. Sci. U S A 1998, 95 (24), 14147–51.

32. Hennig, M.; Geierstanger, B. H., Direct detection of a histidine-histidine side chain hydrogen bond important for folding of apomyoglobin. J. Am. Chem. Soc. 1999, 121 (22), 5123–5126.

33. Andreas, L. B.; Eddy, M. T.; Pielak, R. M.; Chou, J.; Griffin, R. G., Magic angle spinning NMR investigation of influenza A M2(18-60): support for an allosteric mechanism of inhibition. J. Am. Chem. Soc. 2010, 132 (32), 10958–60.

34. Yan, S. H.; Bu, Y. X., Alteration of imidazole dimer on oxidation or water ligation. J. Phys. Chem. B 2004, 108 (36), 13874–13881.

35. Dong, H.; Yi, M.; Cross, T. A.; Zhou, H. X., Ab initio calculations and validation of the pH-dependent structures of the His37-Trp41 quartet, the heart of acid activation and proton conductance in the M2 protein of Influenza A virus. Chem. Sci. 2013, 4 (7), 2776–2787.

36. Morris, G. A.; Freeman, R., Enhancement of Nuclear Magnetic-Resonance Signals by Polarization Transfer. J. Am. Chem. Soc. 1979, 101 (3), 760–762.

37. Hu, J.; Fu, R.; Cross, T. A., The chemical and dynamical influence of the anti-viral drug amantadine on the M2 proton channel transmembrane domain. Biophys. J. 2007, 93 (1), 276–83.

38. Cady, S. D.; Hong, M., Amantadine-induced conformational and dynamical changes of the influenza M2 transmembrane proton channel. Proc. Natl. Acad. Sci. U S A 2008, 105 (5), 1483–8.

