## Supplemental Figures 1 and 2 for "Imidazole-imidazole hydrogen bonding in the pH sensing Histidine sidechains of Influenza A M2"

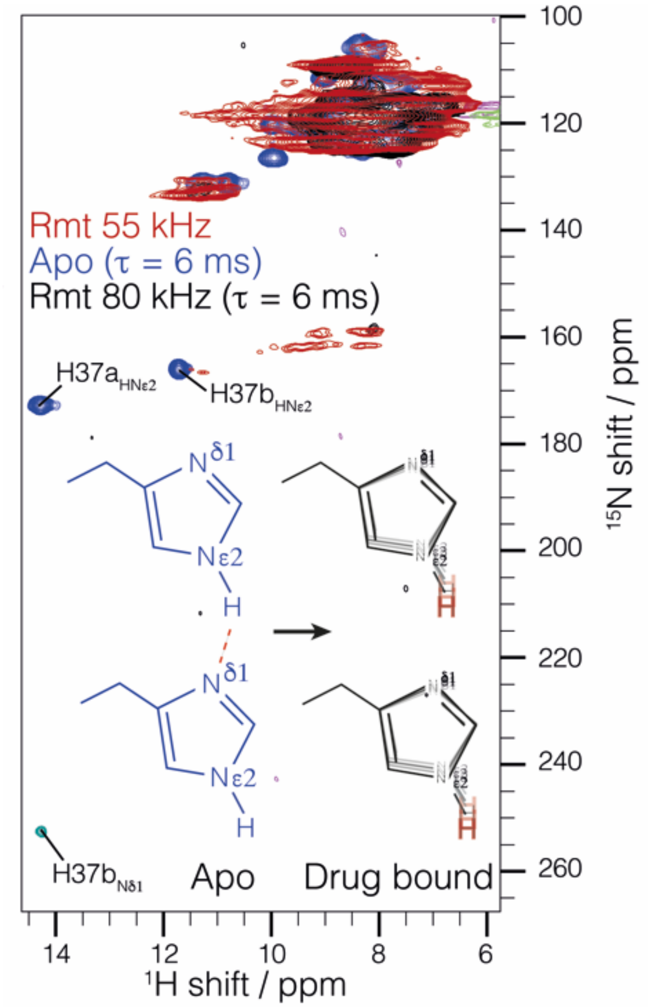


**Figure S1.** Proton detected $^{15}N^{1}H$ correlation spectra indicate destabilization of the imidazole – imidazole hydrogen bond in the presence of the drug Rimantadine. The (H)NH spectrum of M2 in DPhPC lipid bilayers with Rimantadine is shown in red. The proton detected nitrogen INEPT sequence used for J coupling measurement (see Figure 1) is shown in blue (apo) and black (with Rmt). The drug-bound spectrum was recorded in a standard bore 800 MHz Bruker spectrometer using a Bruker 1.3 mm HCN probe with 55 kHz MAS and a thermocouple temperature set at 232K (sample temperature ~15-20 °C). The other spectra were recorded at the 950 MHz Bruker spectrometer with the thermocouple temperature set at 260K (sample temperature ~20 °C). The drug bound sample was recorded with 80 kHz MAS and the apo sample at 100 kHz MAS. Both of these spectra were acquired with an INEPT transfer time τ of 6 ms.

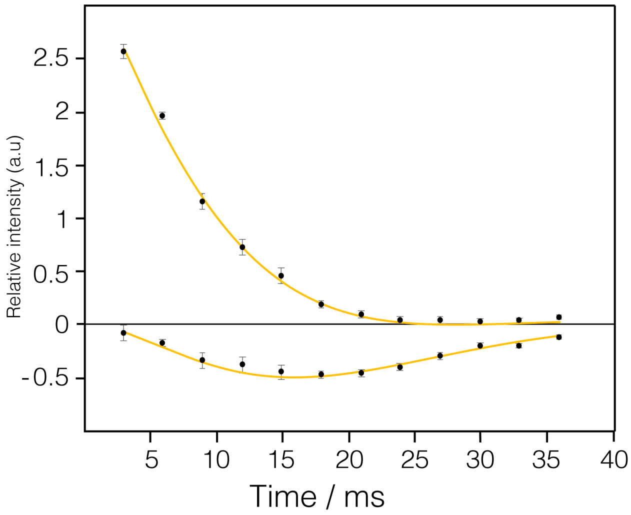


**Figure S2.** Fit of the starting signal on $N_{\varepsilon 2}$ (positive curve) and the buildup of the antiphase term detected at the chemical shift of $N_{\delta 1}$ (negative curve). The curve was fit with the 8.9 Hz J coupling, and a T2 of relaxation time of 44 ms applied to both curves.
